# Avoiding biased estimates: measuring observer avoidance bias in distance sampling estimates of animal populations

**DOI:** 10.1101/2025.09.23.678025

**Authors:** DWE Sankey, CDA Blasi-Foglietti, J Mistry, SK Papworth

## Abstract

Accurate monitoring of animal populations provides crucial information to assess anthropogenic impacts and the success of mitigation efforts. Although distance sampling is the most common method for estimating population densities in mammals and birds, these and other species often violate assumptions that animals do not respond to human observers, causing biased estimates. We present an agent-based model for estimating this bias and evaluating its impact on population density estimates. Animals in our 2D agent-based model follow programmed rules. Models parameterised with either: no human avoidance, or detection of the observer by animals and fleeing behaviours, and are compared to determine the extent of bias that avoidance behaviour can introduce into population density estimates. We test this method with empirical data on responsive movement by Diana monkeys and lesser spot-nosed monkeys from hunted regions of the Gola Forest in Liberia and Sierra Leone. We find that our empirically observed i) distance at which monkeys detected observers and ii) flight initiation distances caused up to a 16% underestimate in population density. We also parameterise the model with theoretical avoidance behaviour to demonstrate the potential scale of the problem, finding up to a 93% density underestimate in extreme – but biologically plausible – cases. By making informed decisions regarding the parameterization of the model, our approach has the potential to complement existing distance sampling techniques, leading to more accurate estimates of species density. This, in turn, can aid in the effective allocation of conservation resources, making our model and its future improvements a valuable tool for researchers in the field.

## INTRODUCTION

Biodiversity worldwide is threatened by climate change, loss and fragmentation of habitats, human disturbance and hunting pressure (Maxwell et al. 2016). To reduce biodiversity loss (Leclère et al. 2020) and assess global progress towards the post-2020 Global Biodiversity Framework under the Convention of Biological Diversity (CBD 2021), accurate monitoring of species numbers is paramount. The IUCN (International Union for Conservation of Nature) Red List (IUCN 2021), relies on estimates of population size to classify species into threat categories, from Least Concern to Critically Endangered. These in turn inform the Red List Index and help guide policies, target conservation measures and inform conventions such as the CBD (Butchart et al. 2005; Betts et al. 2020; IUCN 2021). Likewise, accurate information on the effect of conservation interventions on target populations can aid their evaluation (Stem et al. 2005).

Distance sampling (Buckland et al. 2001) is probably the most widely used method currently adopted to estimate population densities, particularly for mammals (Stokes et al. 2010; Endo et al. 2010; Emerson et al. 2019; Elenga et al. 2020) and birds (Marsden 1999; Magige et al. 2009; Suwanrat et al. 2015). Unlike survey techniques such as plot or strip sampling, this method accounts for uncertain detection scenarios, where not all study objects (individuals/groups/signs) within the survey area are detected (Buckland et al. 2001; Miller et al. 2013). Distance sampling incorporates detection probability in modelled predictions of density using the distance between the transect line/point and the study object. A detection function is fitted to these distances, with probability of detection usually decreasing with increasing distance (Thomas et al. 2010). Key distance sampling assumptions are that 1. *Objects on the line or point are detected with certainty*, 2. *Objects do not move before detection by observers* and 3. *Measurements are exact* (Thomas et al. 2010). These assumptions, however, are unlikely to always be met, particularly when surveying human-avoidant species that perceive humans as a threat (Elenga et al. 2020).

Human-avoidant behaviour is common in many wild species (Whittaker and Knight 1998; Goumas et al. 2020), especially those that are negatively affected by humans (Croes et al. 2006; Tuomainen and Candolin 2011; Patten and Burger 2018). This behaviour is likely greatest for hunted populations, since the cost of not responding may lead to death (Laundré et al. 2010). For example, impala (*Aepyceros melampus*) and greater kudu (*Tragelaphus strepsiceros*) flee at more distant human approaches (higher flight initiation distances – FID) in hunted areas compared to a protected area in Zimbabwe (Tarakini et al. 2014). This responsive movement can lead to increased detections of animals away from their initial location (violating assumption 2), and may mean some animals are not counted, if they move away before they are detected and counted by an observer. Whilst failure to detect an individual may be accounted for by the detection function, together, these behavioural changes may lead to negative detection biases and generate density estimates which are lower than actual densities (Buckland et al. 2001; Elenga et al. 2020).

These lower density estimates may then lead human-avoidant species and populations to be classified as more at risk than they are, diverting limited conservation resources from other species. Furthermore, if these biases have a big enough effect, comparisons between populations of the same species with differences in human-avoidance behaviour could incorrectly lead researchers to conclude there are differences in population density when there are not, affecting evaluation of both anthropogenic effects and conservation interventions (Elenga et al. 2020). This was practically demonstrated by Lindfield et al. (2014) in a study of fished areas and marine protected areas with limited fishing in Guam. As fishing pressure increases, fish avoid the sound of the bubbles produced by standard open SCUBA, negatively biasing density estimates in fished areas. Due to this responsive movement, surveys using open SCUBA found greater differences between the fished areas and marine protected areas than those using non-bubble producing closed-circuit rebreathers (Lindfield et al. 2014). If the standard open SCUBA results had been used to make management recommendations, the effectiveness of the marine protected areas may have been overvalued, or greater than necessary fishing restrictions placed on local communities.

Methods to account for both avoidant and attractant responsive moment in distance sampling have mostly been developed in marine and multi-observer contexts (e.g. Buckland and Turnock 1992; Palka and Hammond 2001), and include searching further ahead along the line, waiting for animals to resume normal behaviours, and double-observer methods (Buckland et al. 2005). However, these methods are not transferable to contexts such as sampling along precut lines in dense jungle (Borchers and Cox 2017). Borchers and Cox (2017) suggested including forward distance (distance between the observer and the location on the transect perpendicular to the animal) as a method to correct for responsive movement. Using case studies on avoidant monkeys and attracted dolphins, they used data simulation to show this approach could reduce bias and increase the probability that 95% confidence intervals will include true population density. This approach was further developed by Elenga et al. (2020) who used the method to account for avoidant responsive movements in blue duikers, but used simulated rather than measured forward distances. Simulations used to quantify bias in both of these studies were based on observed transect-animal distances after avoidant responsive movement, and did not quantify avoidance behaviour. No research has yet quantified the effect of spatially and temporally explicit human avoidant behaviours on density estimates. Here, we achieve this using agent-based models parameterised with both empirical and theoretical data on animal behaviour. This approach contrasts with previous research (Elenga et al. 2020; Borchers and Cox, 2016), which has modelled the effect of responsive movement on observer detection probabilities, but did not explicitly separate how different behaviours which contribute to responsive movement might generate the observed patterns.

Agent-based (also known as individual-based) models offer a bottom-up approach in which outputs emerge from interactions amongst ‘agents’ who behave according to a set of assigned rules (Railsback and Grimm 2019) (for example, human-avoidant movement rules). Agent-based models ultimately aim to study how system level properties come about, and are increasingly being applied to model social and biological systems (DeAngelis and Grimm 2014). They have been used, for example, to predict antipredator behaviour of schooling fish (Vabø and Nøttestad 1997), to estimate collision risks of predatory birds with wind turbines (Eichhorn et al. 2012), and to simulate spread of knowledge of ranger patrol presence amongst hunting communities (Dobson et al. 2019).

Here, we use an agent-based modelling approach to quantify the impact of human avoidant behaviours on density estimates. Firstly, we use novel empirical field data on *i*) distance at which monkeys detected observers and *ii*) flight initiation distances (FID – observer-to-animal distance at which animals fled) from Diana monkeys, *Cercopithecus diana*, and lesser spot-nosed monkeys, *Cercopithecus petaurista* (hereafter referred to as spot-nosed monkeys). These species were studied in both highly hunted and less hunted regions of the Gola region of Liberia and Sierra Leone. Non-human primates (referred to hereafter as primates) were selected as an example as they are particularly vulnerable to anthropogenic disturbances (Kalbitzer and Chapman 2018), and there are documented effects of behavioural changes in hunted populations (Bshary 2001; Hicks et al. 2013). Distance sampling is commonly used to generate density estimates for primates (Buckland et al. 2010), and an estimated 75% of primate species have declining populations due to increasing pressures, such as hunting (Estrada et al. 2017). Second, we modelled theoretical animal populations, with a wide range of human-avoidant behaviours from slightly to highly averse in terms of their *i*) distance at which they detected observers, *ii*) flight initiation distances and *iii*) fleeing distances. As such, we hoped to capture a large spectrum of animal behaviour, enabling researchers to apply our results to a broader range of species.

## METHODS

### Empirical data collection

One set of model simulations were based on empirical data collected in the Gola region of Liberia and Sierra Leone in West Africa. The region is subject to varying levels of hunting pressure across different management areas (see *Supplemental material*). Behavioural data on two West African monkey species was collected between October 2017 and May 2018 along transect lines of 2-4km (see *Supplemental Material*). Diana monkeys, *Cercopithecus diana*, and lesser spot-nosed monkeys, *Cercopithecus petaurista*, were chosen given their high exposure and vulnerability to hunting pressure in West Africa (McGraw et al. 2007; Covey and Mcgraw 2014) and because they can be found across the Gola area (pilot survey May-June 2017). Furthermore, as these species were encountered more often compared to others, larger sample sizes allowed for better informed model parameterisation.

Behavioural variables used for model parameterisation were *i*) the recorded distance at which monkeys detected observers (used to parameterise probability detection curves), and *ii*) the flight initiation distance (FID). After the observers had detected a monkey group, they waited 5 minutes in the same location to determine whether they were detected by the group, which happened on most occasions (Blasi Foglietti 2020). If the observers had not been detected at the end of the 5-minute period, the observer walked towards the group and recorded the distance at which the primate group detected the observer as the detection distance. Detection curves were parameterised so that the median detection distance fell at 0.5 probability (see *Supplemental material*). Following detection, the observers stayed stationary and the group behaviour was observed and recorded for up to a maximum of 20 minutes, providing the group did not flee. Following this period, the observer approached the group further and recorded flight initiation distance (FID) of the closest individual. GPS coordinates for the estimated centre of each group were recorded. The observers then continued walking the transect until the next group was detected. All transects were surveyed following this procedure.

In the model simulation two hunting scenarios were compared to a model parameterised with no avoidance: 1) A lower hunting scenario, parameterised with data collected in the Gola Rainforest National Park (GRNP) in Sierra Leone, a protected area with established on the ground law enforcement in the form of regular patrolling; 2) A higher hunting scenario, parameterised with data collected across the border in Liberia, where a community forest (CF) had no enforced hunting regulations at the time of data collection (see *Supplemental material-Field site* for more details).

### Agent-based model

#### Model set-up

To test the impact of human-avoidant behaviours on density estimates, we set up an agent-based model with “animal groups” responsive to the position of an “observer”. The model represented a two-dimensional virtual world measuring the length of the transect *D_y_*and a border distance *D_border_* around the transect in all directions; Fig. 1). Using a step-length of 25m (Table 1) at each step, the observer sampled any animal groups it could detect (see below in *Distance sampling*) and recorded their perpendicular distance to the transect line. These animal groups were initially positioned across the terrain, at random, using a uniform distribution of x and y axis values (Fig. 1). Animal group starting density was 2.5 groups/km^2^, based on monkey densities recorded at the study site by Klop et al. (2008). At each time step, animal groups that *did not detect and respond to the observer* (Fig. 2A, B), hereafter referred to as normal movement, moved in a random direction (Fig. 2B) at a distance sampled from an exponential decay distribution (Fig. 2A). Here, the most likely distance moved is zero (stationary), and greater distances become less and less likely. This behaviour resembles monkey group movements around a home range (Oates 2011).

**Figure 1.**
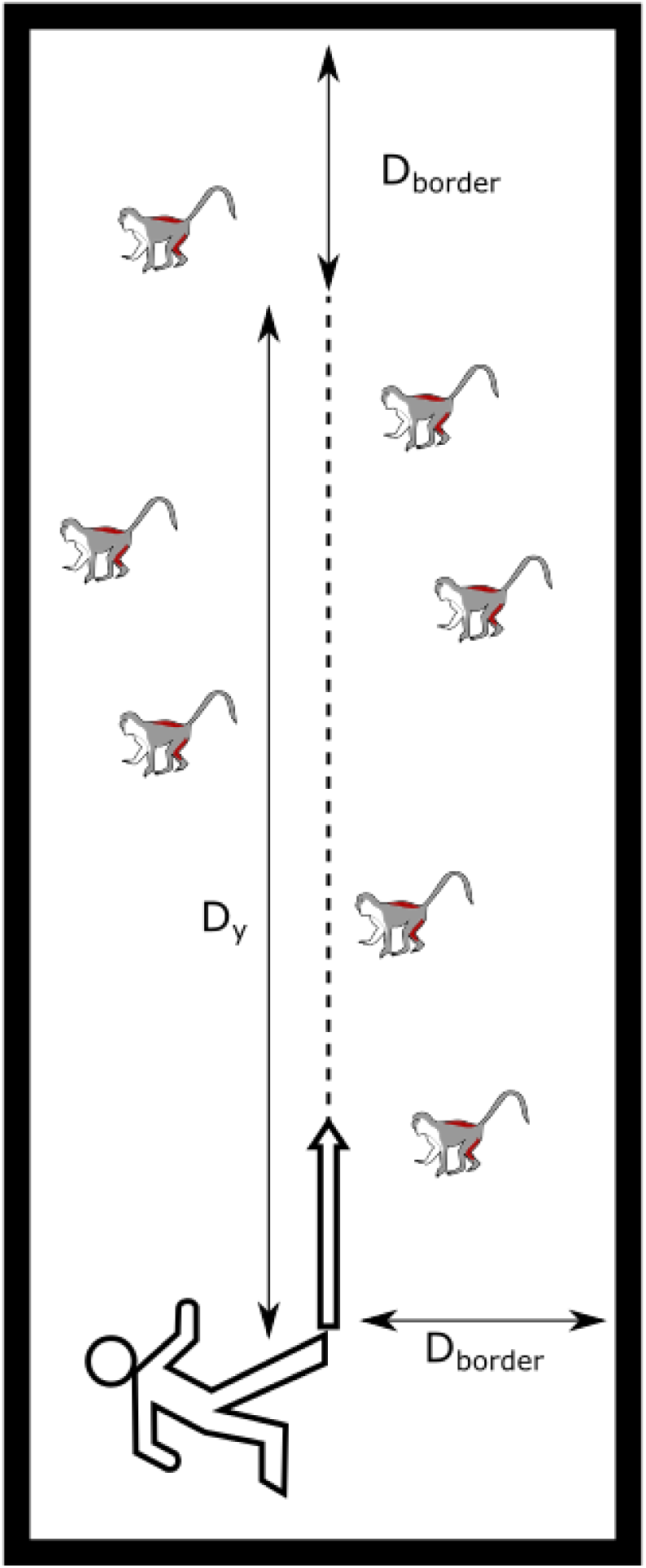
Schematic representation of the spatial landscape of the model showing the length of transect lines *D_y_*, the transect to border distance *D_border_*, the observer starting location and the uniformly distributed animal groups (represented by a single monkey).

**Figure 2.**
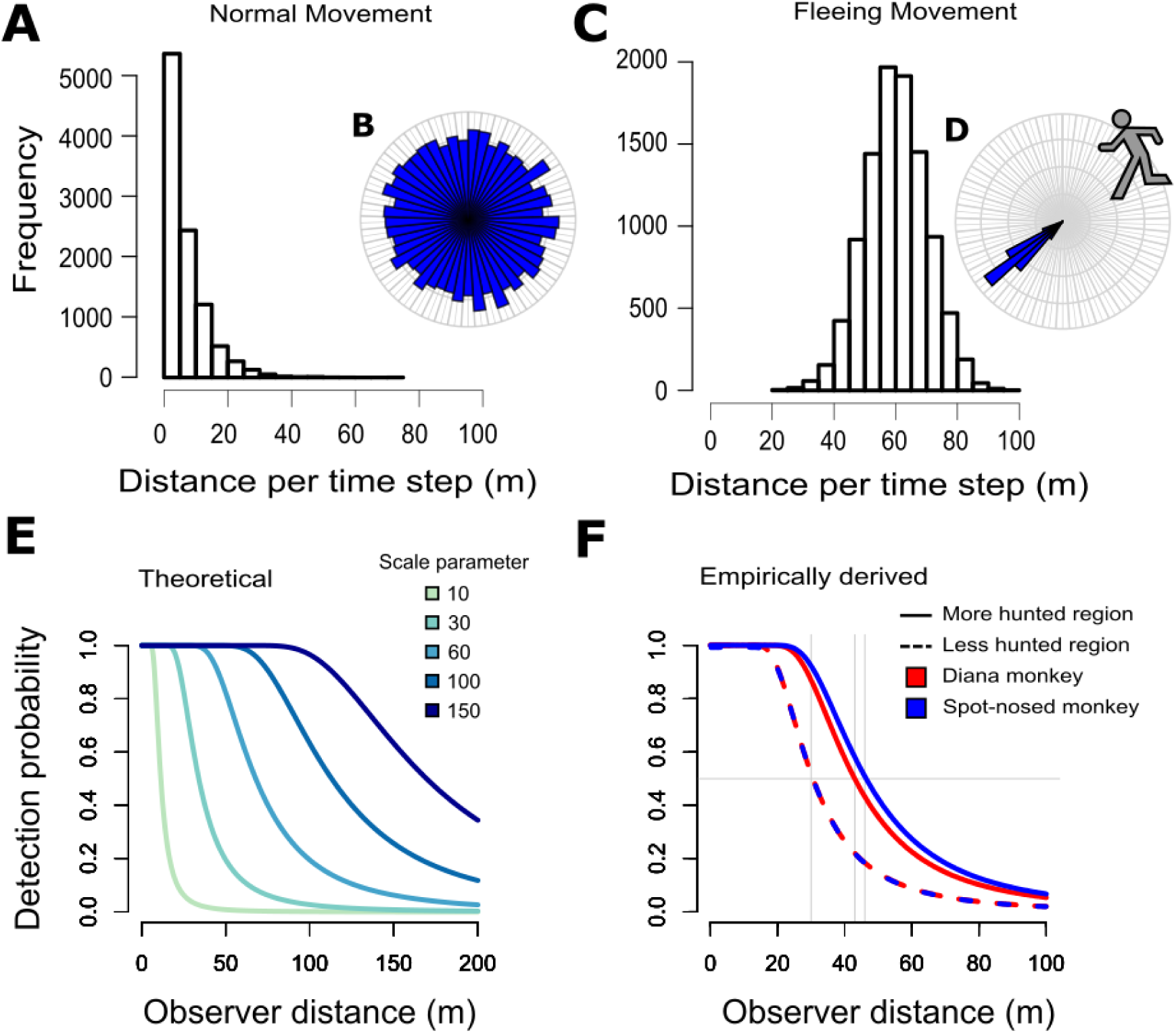
Parameterisation for all models. Normal animal group movement at each time step was **A** mostly slow, with high speeds being rare (rate of exponential decay = 0.15), and **B** direction of movement was a random walk from –π to π radians. In fleeing groups, **C** larger distances are covered per time step (60m mean; 15m standard deviation, see Table 1 for values in theoretically parameterised models), with a **D** highly skewed distribution for direction of movement away from the observer (Von-Mises distribution with *k* = 50). **E** Theoretical detection distances (scale parameters) for animal groups (see legend) produced detection probability curves as shown (coloured lines). **F** Detection distances from empirical data were fitted so that the median observed distance at which monkeys spotted human observers (vertical grey lines) produced a detection probability of 0.5 (horizontal grey line) (See *Supplemental material*).

**Table 1.**
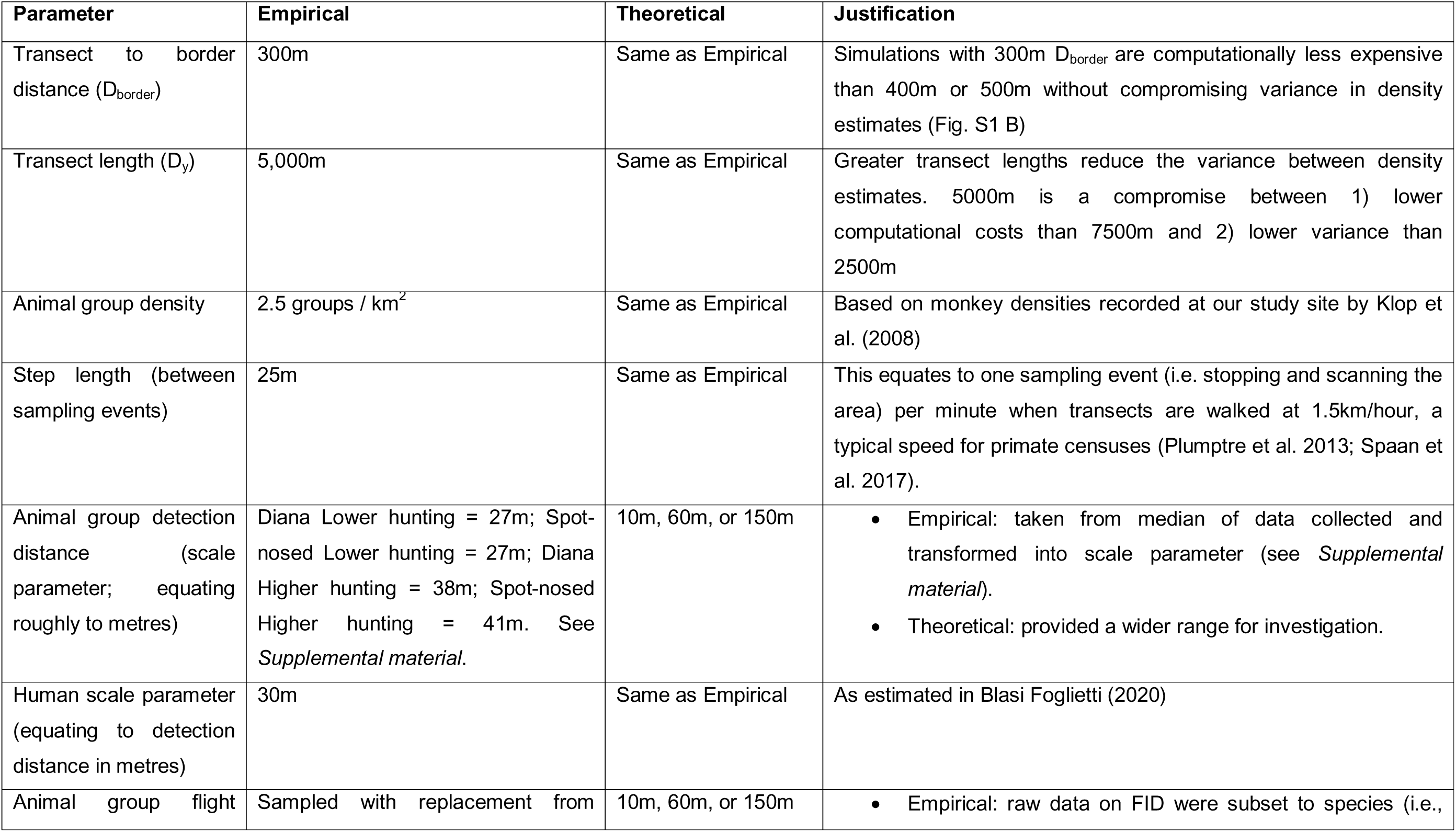
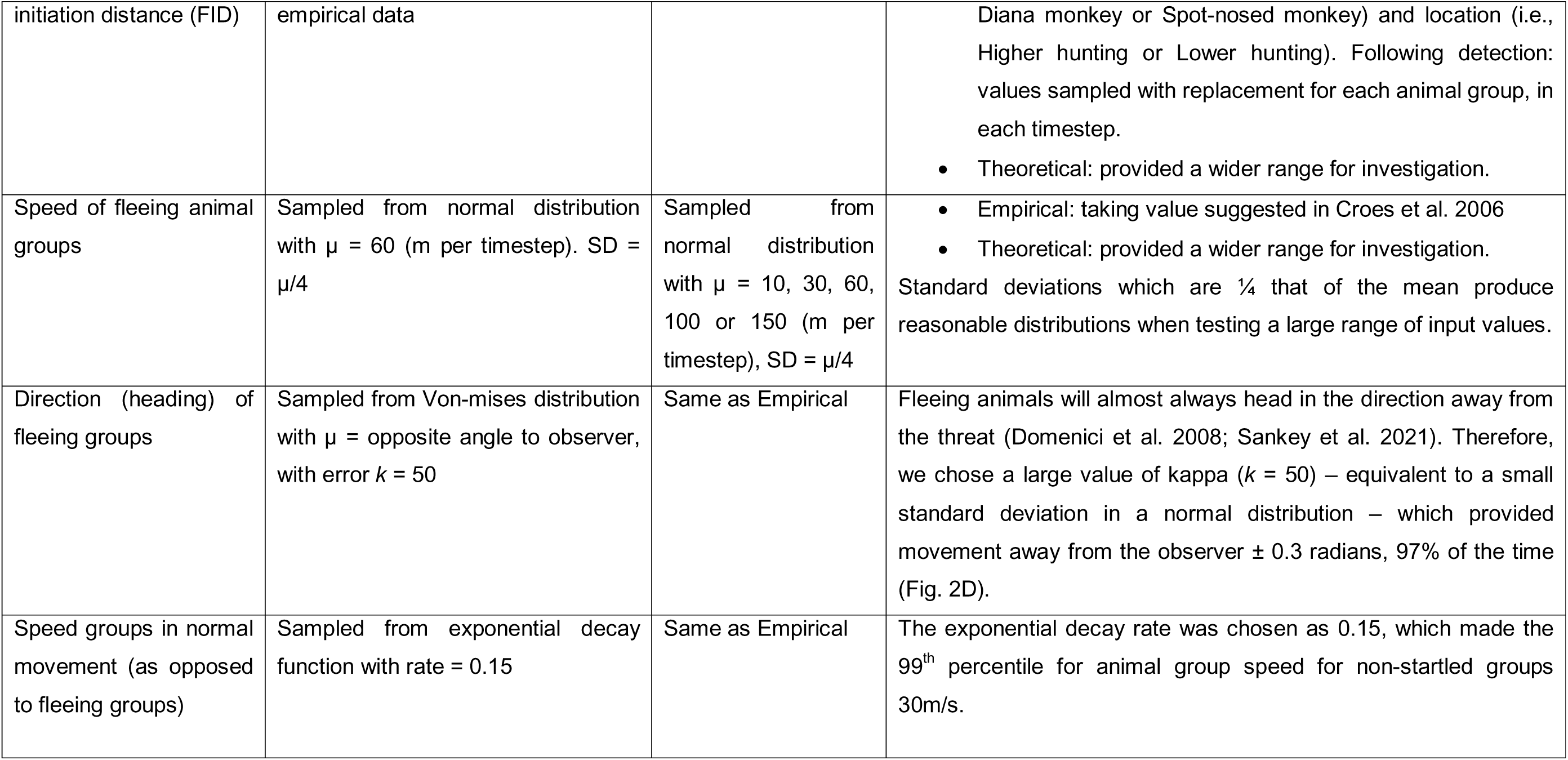
Agent-based model parameters across our two model types (empirical data driven and theoretical).

#### Avoidance behaviour

The other state a group can take is fleeing, which requires two conditions: (i) the observer must be detected, and (ii) the observer must be within the animal group’s allocated flight initiation distance (FID) (see Fig. 2C-F). Firstly, the likelihood of an animal detecting the observer was (1) for empirically parameterised models: derived from real data of monkey detection distances (see Fig. 2F and *Supplemental material*), or (2) for theoretical simulations: given a scale parameter of 10m, 30m, 60m, 100m or 150m (Fig. 2E). If the first condition was met (observer was spotted), animal groups were then programmed to flee or not based on whether distance to the observer was closer than the group’s allocated FID. For empirically parameterised models, FID was sampled with replacement from empirical observations of FID in each time step (see data in Results and Discussion). For theoretical simulations FID was given consistent values of 10m, 30m, 60m, 100m or 150m.

For fleeing groups, movement was faster (Fig. 2C) and directional: away from the observer (Fig. 2D). We chose fleeing direction from a von Mises distribution (a circular version of a normal distribution; Agostinelli and Lund 2013), parameterised so that movement was directly away from the observer (± 0.3 radians) 97% of the time (Table 1). The distance that fleeing animals travelled in one time-step was sampled from a normal distribution. For empirically driven simulations mean fleeing distance was 60m, which was parameterised from observed data of multiple Cercopithecus monkey species fleeing distances (Croes et al. 2006 who noted distances of “typically >50 m”). For theoretical simulations fleeing distance was given mean values of 10m, 30m, 60m, 100m, or 150m per time step. Standard deviation of fleeing distance was always equal to the mean value divided by four, for consistency, and to reduce the chance of negative distances for smaller mean values. Dividing by four, as opposed to any other number, is largely arbitrary and worthy of further attention in future work.

#### Distance sampling

Just as the animal groups could detect the observer; the observer could also detect the animal groups. The further the animal group was from the observer, the less likely the observer was to detect the group, with a probability following a hazard-rate detection curve (e.g., Fig. 2 E,F). We parameterised the detection curve for humans using a median of the values we observed in the animal groups, as monkey groups in our field study tended to detect humans at similar distances to the distance that humans detected the monkey groups (Blasi Foglietti 2020). When detected, the observed animal group’s perpendicular distance to the transect line was recorded, and the group was deleted from the model to prevent multiple observations of the same group. The distribution of perpendicular distances (Buckland et al. 2001) was used to fit the detectability curve to produce an overall density estimate (using “*Distance*” package in R; Miller et al. 2016).

All the previously mentioned actions take place in each time step of the model. The animal groups then move towards their new location (after either a fleeing or normal movement step), and the observer moves into the new position, 25m along the transect line.

We calculated 1000 density estimates for each main or supplemental analysis reported in this paper, each of which was calculated from 100 simulated transects. Thus, we had N=1000 density estimates for each set of model conditions. We do not report confidence intervals around the median density estimates from our simulations (White et al. 2014). This is due to the relationship between model parameters and precision of estimates. As an example, larger transect lengths would decrease the reported variance (Fig. S1), as would increasing the number of transects used in each density estimate calculation.

### Ethical Statement

All fieldwork was conducted in accordance with national and institutional guidelines for the ethical treatment of wildlife. Research protocols were approved by Royal Holloway University of London Ethics Committee and carried out under research permits issued by the relevant authorities in Sierra Leone and Liberia. Observers followed established best-practice guidelines for minimising disturbance to wild primates. No animals were handled, captured, or otherwise physically disturbed during this study.

## RESULTS AND DISCUSSION

Consistent with being more human-averse, monkeys from the higher hunted region demonstrated increased *i*) distance at which they detected observers and *ii*) flight initiation distances than those from the less hunted region (linear model with hunting region and species as additive covariates: t = –3.311, DF = 53, p = 0.002; flight initiation distance – LM: t = –3.359, DF = 53, p = 0.001; Fig. 3). With these greater distances at which they detected observers and flight initiation distances, the animal groups within the higher hunting scenario of the agent-based model were more likely to detect the observer and then respond (by fleeing) before they were detected by the observer. This breaks the distance sampling assumption that “Objects do not move” before being detected. This movement, in turn, meant that animal groups were less likely to be detected by the observer in the subsequent time-step. We found density estimates decreased by 12.0% (Diana monkeys) and 16.1% (Spot-nosed monkeys) when comparing models parametrised with empirical data from higher hunted regions to models programmed with no avoidance (Fig. 4). Models using parameters from lower hunted regions showed a more modest decrease, by 2.9% (Diana monkeys) and 2.5% (Spot-nosed monkeys). Note that median estimated densities from the model were 2.82 groups/km^2^, higher than the simulated input measurement of 2.5 groups/km^2^ (Fig. S1). This may be due to documented effects of animal movement on distance sampling estimates – as animal speed increases, the number of detections near the line increase, positively biasing estimates (Glennie et al. 2015). We therefore report our results relative to our baseline, no-avoidance, values.

**Figure 3.**
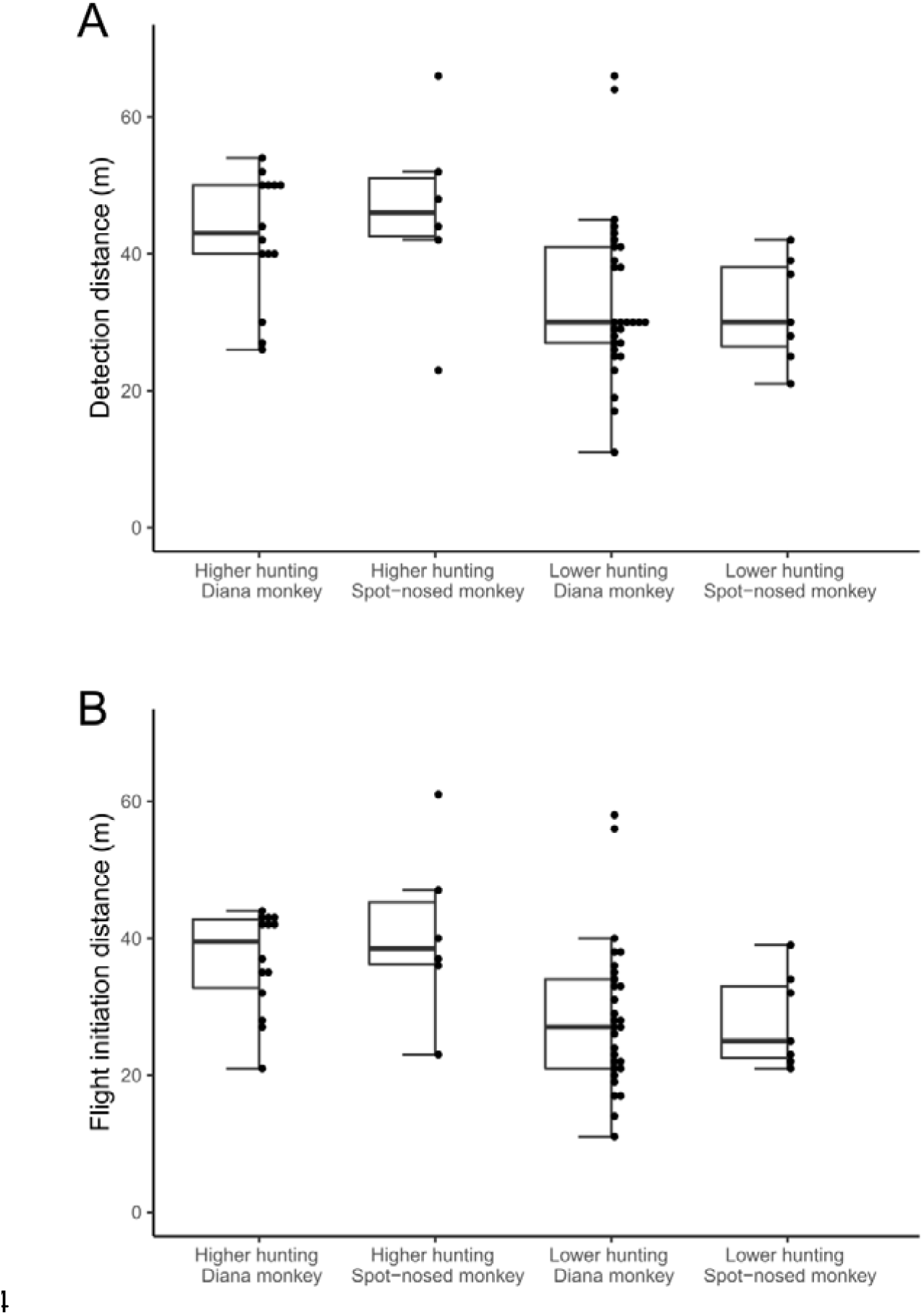
Empirical data. **A**) Distance (m) at which monkeys detect the human observer and **B)** distance at which they fled (FID) following an approach by the observer across our conditions.

**Figure 4.**
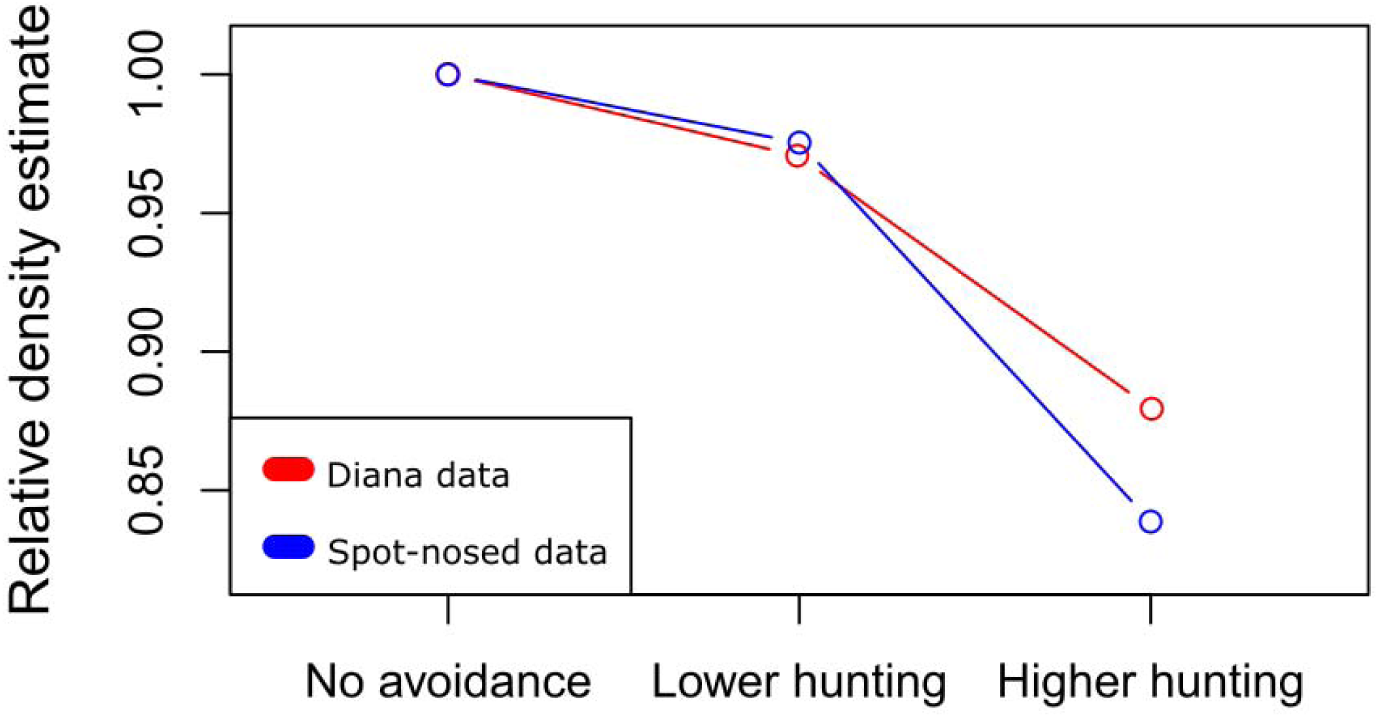
Models parameterised with empirical data. Relative density estimate: median of N=1000 density estimates (animal groups per square km) relative to the model parameterised with no avoidance (left). Real data collected on distance at which monkeys detect observers and flight initiation distances from Diana (red) and lesser spot-nosed monkeys (blue) from both a less hunted region and a more heavily hunted region were programmed into our model to produce “Lower hunting” and “Higher hunting” estimates. The model assumed an actual population density 2.5 animal groups per km^2^.

Based on the results of this analysis, we estimate that in higher hunted regions of the Gola Forest the actual density would be between 13.6-19.1% higher than estimates generated using distance sampling. However, the density estimates in our lower hunting scenario are relatively closer to the actual values (Figure 4), consistent with the expectation that stronger behavioural reactions to humans will introduce greater bias into density estimates (Elenga et al. 2020).These differences with hunting pressure may be observable in monkeys due to their plastic responses to humans (Papworth et al. 2013; Sheehan and Papworth 2019; LaBarge et al. 2020). In species which are always human-averse (Frid and Dill 2002), more consistent responses to humans may mean that the potential bias as a result of avoidant behaviour is the same regardless of hunting pressure.

Next, we parameterised the model with theoretical parameters, to explore the impact of different aspects of avoidance behaviour on distance sampling. We reveal that– *i*) flight initiation distance, *ii*) fleeing distance and *iii*) distance at which monkeys detected observers– can all drive bias in distance sampling methods, with all three reducing the density estimate. This decrease is sharp at first, but plateaus for larger values of these variables (Fig. 5).

**Figure 5.**
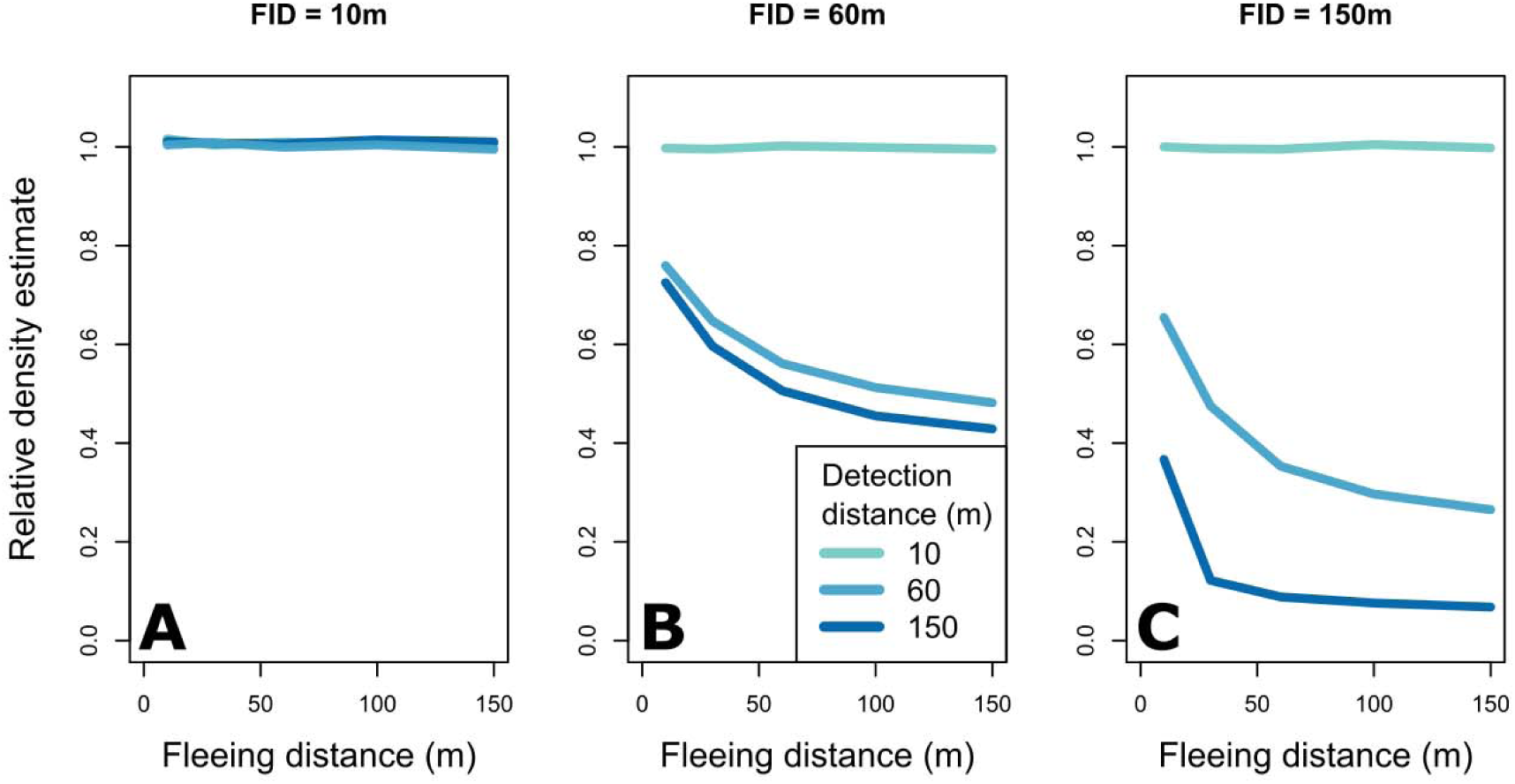
Models with theoretical parameters. Relative density estimate (see below) is plotted against fleeing distance, when flight initiation distance (FID) was either **A** 10m, **B** 60m, or **C** 150m. Coloured lines indicate parameterised distance (m) at which animals detect observers as seen in legend. Relative density estimate: median of N=1000 density estimates (animal groups per square km) relative to the model parameterised with no avoidance.

We show density estimates more than 90% lower than the density for the simulations with no avoidance behaviour (Fig. 5, panel C), when distance at which monkey detect observers and FID distance are 150m, and fleeing distances are over 60m. Based on the most extreme values used in the parameterisation of our model, *actual densities* of animals in the wild could be up to ten times higher than their *estimated densities*. The parameters which produced an 90% reduction in density estimate are not unfeasible. 150m detection range of animals (and hence flight initiation distance too) is not excessive, given, for example, elephants capabilities to sense over several kilometres (Römer 2001). Fleeing distances which led to over 90% bias were over 60m per timestep –twice the observer speed of 25m/timestep. Animal movement more than twice the speed of humans is both common (Ruf et al. 2006), and a known source of bias in distance sampling studies (Glennie et al. 2015), even when this movement is not in response to observers. Together, this information can help identify species and individuals whose responses to humans are most likely to bias density estimates. Our results suggest that the potential for bias is relatively minor when either FID or distance at which animals detect observers is 10m, most likely as the probability of detection by observers in our model is high up to 30m, so animals are detected before they are flushed. In contrast, when distance at which animals detect observers distances and FID are greater than observer detection distances, even if animals only flee a short distance (e.g. 10m, Fig. 5), biases can be substantial. A rich body of research is available on FID for human disturbances which can identify which animals are most likely to be effected by these biases (Dill and Frid 2020).

### Using our methods to estimate distance bias

As distance sampling is a common density estimation method (Stokes et al. 2010; Endo et al. 2010; Emerson et al. 2019; Elenga et al. 2020; Marsden 1999; Magige et al. 2009; Suwanrat et al. 2015), which feeds into population estimates that determine IUCN red list status (Butchart et al. 2005; Betts et al. 2020; IUCN 2021). This in turn can inform allocation of resources to threatened species. We encourage empiricists who use distance sampling methods, and who can also collect avoidance behaviour data, to account for distance sampling biases. This can be done by running simulations on our web application available at (webpage currentlyu and running in RStudio. Data collection for avoidance behaviour can be run in tandem with distance sampling data collection. Flight initiation distances, distances at which observers were spotted, and perpendicular distances to the transect, are all distance measures and thus estimation of all three parameters requires little extra effort than just one of these metrics. Additional information, such as distance along the line could also be collected. Distance along the line can be used to reduce biases introduced by movement in response to observers (Bouchers and Cox, 2017). At present however, the R package for this requires customization (see Elenga et al. 2020) and so cannot currently be incorporated into this web application for general use.

### Extending our model to account for other human avoidant behaviour

Fleeing and heightened vigilance are not the only observed human avoidant behaviours documented in the wild. Animals may reduce vocalisations around humans (Bshary 2001; Hicks et al. 2013), display freezing behaviours (Kümpel et al. 2008), change the size of the groups they associate with (Watanabe 1981; Dooley and Judge 2015), or make temporal shifts in activity to periods of human absence (Ohashi et al. 2013, Marchand et al. 2014). We provide a table of these behaviours, and how they might be implemented in future models (Table 2). Meticulous parameterisation of how, for example, freezing behaviour actually decreases detectability, could support real world applicability.

**Table 2.**
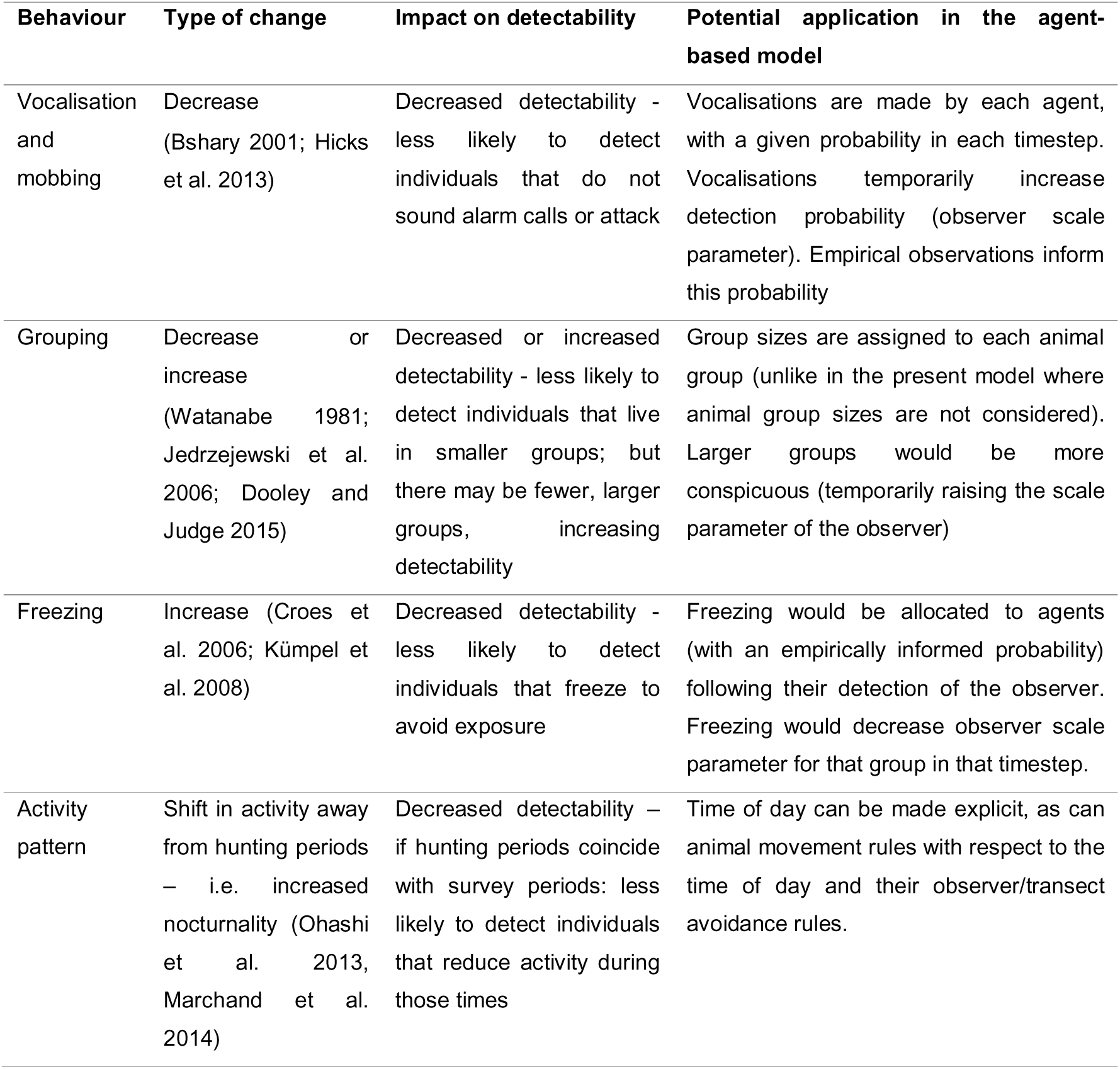
Examples of how to apply other behavioural shifts due to anthropogenic pressure to our agent-based model.

### Conclusion

We provide a method to estimate the bias in distance sampling methods when applied to human avoidant animals. Quantifying this bias can enable researchers to understand their estimates of species density more accurately, which in turn can inform an effective conservation strategy. We trial the method using data collected on Diana monkeys and Spot-nosed monkeys from the Gola Rainforest National Park and found that density estimates of our study species from higher hunted regions would likely be underestimates.

Furthermore, we demonstrate that even larger biases are possible, as under theoretical parameterisation species densities estimates were as low as 7% of their actual densities for biologically plausible avoidance metrics. These results underscore the importance of accounting for avoidance behaviour when estimating species density. Overall, our method has the potential to improve the accuracy of red list status assessments for a variety of animal species, provided that avoidance behaviour data is available.

## SUPPLEMENTAL MATERIAL

### Parameterisation of detection curves

Detection functions are an essential part of distance sampling. We chose to use hazard-rate curves, due to the presence of a shoulder where detection is high (see Hayes and Buckland 1983; Buckland et al. 2001). We used an intermediate value of three as the “shape parameter” (see Buckland et al. (2001)); but also this is adjustable in the web application) for all models and adjusted the “scale” parameter to match our empirical data. We allocated a scale parameter which gave a detection probability of 0.5 at the median observed “detection distance” (see: empirical data collection) at which the monkeys spotted the human observer. For example, the median detection distance of human observers by the Diana monkeys in the more hunted region was 43m. The detection curve which most closely matched 43m at the 0.5 intercept was a detection curve with a scale of 38 (see Fig. 2F for all parameterised curves). This was repeated for both monkey species at both field sites.

### Field site

The study region comprised two adjacent areas. The Gola Rainforest National Park (GRNP) of Sierra Leone (7°61’N, 10°94’W) covers an area of 700 km^2^ and was inaugurated in 2011. The park has established on-the-ground law enforcement in the form of regular ranger patrols deployed throughout the park, covering around 7000 km per year (Barca et al. 2018b). The effectiveness of the park’s conservation interventions is highlighted by positive figures showing increasing populations of the Endangered Upper Guinea red colobus, *Piliocolobus badius*, for example (Barca et al. 2018a). This suggests an effective reduction in hunting pressure since the establishment of the park. Data from this area were used to parameterise the lower hunting scenarios within the model.

The second area is a community forest (CF) in Liberia (7°74’N, 10°52’W) which covers about 400 km^2^ adjacent to the northern parts of the GRNP. Through a partnership between the communities, the Government of Liberia and conservation NGOs (the Society for Conservation of Nature in Liberia, SCNL, and the Royal Society for the Protection of Birds, RSPB), the GolaMA project (running between 2014 and 2019) has been working to implement and establish this community forest, with the aim of achieving sustainable management of forest resources outside protected areas (Jones et al. 2018). For simplicity this area will be referred to as the community forest hereafter. Work by Jones et al. (2018) indicates that hunting in the community forest is widespread. Data from this area were used to parameterise the higher hunting scenarios within the model. Data were also collected in the adjacent Gola Rainforest National Park of Liberia which was demarcated in 2018 (Blasi-Foglietti, 2020), but were not used to parametrize the models in this study due to the new management status at the time of data collection and uncertainty about levels of historical and contemporary hunting pressure.

Different gun regulations between countries, with stricter rules in Sierra Leone compared to Liberia, may also translate into different degrees of hunting pressure between areas (Lahai 2013). Therefore, overall differences in both local management and country level legislations likely result in different patterns of biodiversity exploitation and degree of hunting pressure across the Gola forest region, with lower hunting pressure expected in the GRNP of Sierra Leone and higher hunting pressure expected in the community forest of Liberia.

### Behavioural data collection

Behavioural observations were conducted between October 2017 and May 2018. In Sierra Leone data was collected between October and November 2017 and between January and March 2018, in Liberia data was collected between November and December 2017 and between March and May 2018. Alternating between countries helped to account for potential variation linked to time of year.

A total of 18 permanent 4-km transects were monitored (10 in the GRNP, 8 in the community forest). These lines are normally used as part of the ongoing monitoring in both countries. Additionally, 11 2-km lines were cut specifically for this study at least 2 weeks prior to the data collection phase in the GRNP. The 2-km transects and the permanent transects in Sierra Leone were spaced at a distance of 2.5 km, whereas the permanent transects in Liberia were spaced at a distance of 1.5 km. All transects were sampled once to prevent recording responses from the same group twice, with the exception of 3 lines in Liberia, which were sampled a second time as no observations were recorded during the first attempt. Observations on different transect lines are expected to be independent as the home rages of the focus species are smaller than the distance between transect lines. Total survey effort was thus 94 km across the survey region, with 62 km walked in the GRNP and 32 km walked in the community forest.

Observations started in the morning, between 6:45 am and 7:30 am. Each day a 2-km transect was monitored. The 4-km transects were split into 2 and monitored on 2 consecutive days. Survey teams varied between 2-4 people, with CBF present during all observations. Observers walked transects at a slow pace (0.5-1km/h), scanning the surroundings with the use of binoculars and listening for monkey calls. Transects were walked in silence to minimise the chance of being detected by animal groups/individuals. If monkeys were located through vocalisation, the observers left the transect line (up to a maximum of 500 m) towards the direction of the calls and returned to the same point along the transect to continue the survey once the observation was completed.

Upon detection of a primate group (through sight or vocalisation), the experimental phase would begin. The observers remained hidden out of view for a period of 5 minutes (or until detected) to gather baseline group-level information on primate behaviour prior to detection. Species, number of individuals within the group, group cohesion and presence of other species was recorded.

If the observers had not been detected at the end of the 5-minute period, the observer walked towards the group and recorded the distance at which the primate group detected the observer as the detection distance.

Following detection, the group was observed for up to a maximum of 20 minutes, providing the group did not flee. Following this period, the observer approached the group further and recorded flight initiation distance (FID) of the closest individual. GPS coordinates for the estimated centre of each group were then recorded. The observers then continued walking the transect until the next group was detected. All transects were surveyed following this procedure.

All GPS coordinates were collected using a Garmin GPSMAP 64s handheld navigator. Time was kept using a Casio F-91W wristwatch and vocalisations were recorded with a Marantz PMD661 Portable Stereo recorder with directional microphone and a compact Sony handheld recorder.

### Sensitivity analyses

We performed 100000 iterations (with density calculated every 100 transects) of *D_y_*and *D_border_* set at different levels, to gauge the effect of these variables on our results, and also to decide on which would be best as a default.

**Figure S1.**
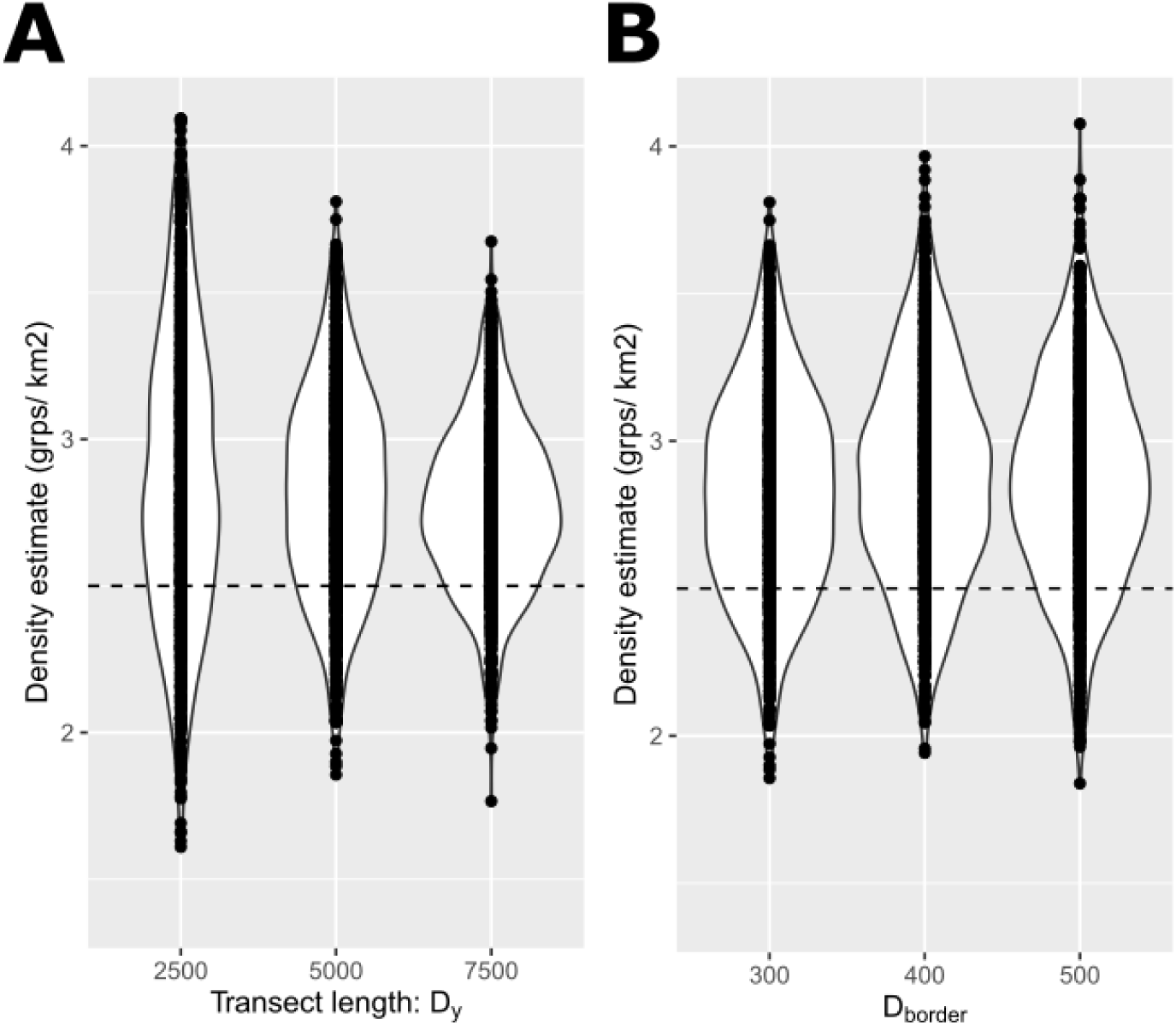
Sensitivity analyses. Each violin plot represents 1000 densities, each calculated from 100 transects in the model. All using conditions with no reaction to the observer (FID parameter was scaled to 0). The estimated densities are generally higher than the input measurement 2.5 groups/km^2^, median = 2.82, representing a 12.8% increase on the input density. This is why results are reported as relative to these baseline values. **A)** an investigation of transect length *D_y_*. Variance between density estimates decreases with increasing transect length. 5000m transect lengths were chosen due to a compromise between faster computation (than 7500m) and lower variance (than 2500m). During these iterations, the value for *D_border_* was always 300. **B)** an investigation of *D_border_*. Iterations with *D_border_*= 300m had faster computations and no observable difference between density estimates compared with 400m and 500m, and so was chosen as a default value for the main results. During these iterations, the value for D_y_ was always 5000.

## REFERENCES

1. Agostinelli C, Lund U (2013) R package circular: Circular Statistics (version 0.4-3). URL https://r-forge r-project org/projects/circular

2. Barca B, Tayleur C, Turay BS (2018a) Recovery of Upper Guinea red colobus Piliocolobus badius and other primates in Gola Rainforest National Park, Eastern Sierra Leone. In: 27th International Primatological Society Congress

3. Barca B, Turay BS, Kanneh BA, Tayleur C (2018b) Nest Ecology and Conservation of Western Chimpanzees (*Pan troglodytes verus*) in Gola Rainforest National Park, Sierra Leone. Primate Conserv 32:133–139. doi: www.primate-sg.org/primate_conservation/

4. Betts J, Young RP, Hilton-Taylor C, Hoffmann M, Rodríguez JP, Stuart SN, Milner-Gulland EJ (2020) A framework for evaluating the impact of the IUCN Red List of threatened species. Conserv Biol 34:632–643. doi: 10.1111/COBI.13454

5. Blasi Foglietti CDA (2020) Hunting pressure and primate behaviour in the Gola forest of West Africa. Royal Holloway University of London

6. Borchers DL, Cox MJ (2017) Distance sampling detection functions: 2D or not 2D? Biometrics 73:593–602

7. Bshary R (2001) Diana monkeys, *Cercopithecus diana*, adjust their anti-predator response behaviour to human hunting strategies. Behav Ecol Sociobiol 50:251–256. doi: 10.1007/s002650100354

8. Buckland ST, Anderson DR, Burnham KP, Laake JL (2005) Distance sampling. Encycl Biostat 2:

9. Buckland ST, Anderson DR, Burnham KP, Laake JL, Borchers DL, Thomas L (2001) Introduction to Distance Sampling, estimating abundance of biological populations. Oxford University Press, Oxford

10. Buckland ST, Plumptre AJ, Thomas L, Rexstad EA (2010) Design and analysis of line transect surveys for primates. Int J Primatol 31:833–847

11. Buckland ST, Turnock BJ (1992) A robust line transect method. Biometrics 901–909

12. Butchart SH., Stattersfield A., Baillie J, Bennun L., Stuart S., Akakaya H., Hilton-Taylor C, Mace G. (2005) Using Red List Indices to measure progress towards the 2010 target and beyond. Philos Trans R Soc B Biol Sci 360:255–268. doi: 10.1098/RSTB.2004.1583

13. CBD (2021) First Detailed Draft of the new Post-2020 Global Biodiversity Framework. Convention on Biological Diversity

14. Covey R, Mcgraw WS (2014) Monkeys in a West African bushmeat market: implications for cercopithecid conservation in eastern Liberia. Trop Conserv Sci 7:115–125. doi: 10.1177/194008291400700103

15. Croes BM, Laurance WF, Lahm S., Tchignoumba L, Alonso A, Lee M., Campbell P, Buij R (2006) The influeunce of hunting on antipredator behaviour in Central African monkeys and duikers. Biotropica 39:257–263. doi: 10.1111/j.1744-7429.2006.00247.x

16. DeAngelis DL, Grimm V (2014) Individual-based models in ecology after four decades. F1000Prime Rep 6:. doi: 10.12703/p6-39

17. Dill LM, Frid A (2020) Behaviourally mediated biases in transect surveys: a predation risk sensitivity approach. Can J Zool 98:697–704

18. Dobson ADM, de Lange E, Keane A, Ibbett H, Milner-Gulland EJ (2019) Integrating models of human behaviour between the individual and population levels to inform conservation interventions. Philos Trans R Soc B Biol Sci 374:20180053. doi: 10.1098/rstb.2018.0053

19. Domenici P, Booth D, Blagburn JM, Bacon JP (2008) Cockroaches keep predators guessing by using preferred escape trajectories. Curr Biol 18:1792–1796

20. Dooley HM, Judge DS (2015) Kloss gibbon (*Hylobates klossii*) behavior facilitates the avoidance of human predation in the Peleonan forest, Siberut Island, Indonesia. Am J Primatol 77:296–308. doi: 10.1002/ajp.22345

21. Eichhorn M, Johst K, Seppelt R, Drechsler M (2012) Model-based estimation of collision risks of predatory birds with wind turbines. Ecol Soc 17:art1. doi: 10.5751/ES-04594-170201

22. Elenga G, Bonenfant C, Péron G (2020) Distance sampling of duikers in the rainforest: Dealing with transect avoidance. PLoS One 15:e0240049. doi: 10.1371/JOURNAL.PONE.0240049

23. Emerson LD, Ballard G-A, Vernes K, Emerson LD, Ballard G-A, Vernes K (2019) Conventional distance sampling versus strip transects and abundance indices for estimating abundance of greater gliders (Petauroides volans) and eastern ringtail possums (Pseudocheirus peregrinus). Wildl Res 46:518–532. doi: 10.1071/WR18155

24. Endo W, Peres CA, Salas E, Mori S, Sanchez-Vega JL, Shepard GH, Pacheco V, Yu DW (2010) Game vertebrate densities in hunted and nonhunted forest sites in Manu National Park, Peru. Biotropica 42:251–261. doi: 10.1111/j.1744-7429.2009.00546.x

25. Estrada A, Garber PA, Rylands AB, et al. (2017) Impending extinction crisis of the world’s primates: Why primates matter. Sci Adv 3:e1600946. doi: 10.1126/sciadv.1600946

26. Frid A, Dill L (2002) Human-caused disturbance stimuli as a form of predation risk. Conserv Ecol 6:

27. Glennie R, Buckland ST, Thomas L (2015) The effect of animal movement on line transect estimates of abundance. PLoS One 10:e0121333

28. Goumas M, Lee VE, Boogert NJ, Kelley LA, Thornton A (2020) The Role of Animal Cognition in Human-Wildlife Interactions. Front. Psychol. 11:3019

29. Hayes RJ, Buckland ST (1983) Radial-distance models for the line-transect method. Biometrics 39:29’42. doi: 10.2307/2530804

30. Hicks TC, Roessingh P, Menken SBJ (2013) Impact of humans on long-distance communication behaviour of Eastern Chimpanzees (*Pan troglodytes schweinfurthii*) in the Northern Democratic Republic of the Congo. Folia Primatol 84:135–156. doi: 10.1159/000350650

31. IUCN (2021) The IUCN Red List of Threatened Species. Version 2021–1

32. Jedrzejewski W, Spaedtke H, Kamler JF, Jedrzejewska B, Stenkewitz U (2006) Group size dynamics of red deer in Białowieża Primeval Forest, Poland. J Wildl Manage 70:1054– 1059

33. Jones S, Keane A, St John F, Vickery J, Papworth S (2018) Audience segmentation to improve targeting of conservation interventions for hunters. Conserv Biol 33:895–905. doi: 10.1111/cobi.13275

34. Kalbitzer U, Chapman CA (2018) Primate responses to changing environments in the Anthropocene. In: Kalbitzer U, Jack K. (eds) Primate life histories, sex roles, and adaptability. Springer nature Switzerland, pp 283–310

35. Klop E, Lindsell J, Siaka A (2008) Biodiversity of Gola Forest, Sierra Leone

36. Kümpel NF, Milner-Gulland EJ, Rowcliffe JM, Cowlishaw G (2008) Impact of gun-hunting on diurnal primates in continental Equatorial Guinea. Int J Primatol 29:1065–1082. doi: 10.1007/s10764-008-9254-9

37. LaBarge LR, Hill RA, Berman CM, Margulis SW, Allan ATL (2020) Anthropogenic influences on primate antipredator behavior and implications for research and conservation. Am J Primatol 82:e23087

38. Lahai K (2013) The distribution of key species in the Gola Forests

39. Laundré JW, Hernández L, Ripple WJ (2010) The landscape of fear: ecological implications of being afraid. Open Ecol J 3:1–7

40. Leclère D, Obersteiner M, Barrett M, et al. (2020) Bending the curve of terrestrial biodiversity needs an integrated strategy. Nat 2020 5857826 585:551–556. doi: 10.1038/s41586-020-2705-y

41. Lindfield SJ, Harvey ES, McIlwain JL, Halford AR (2014) Silent fish surveys: bubble-free diving highlights inaccuracies associated with SCUBA-based surveys in heavily fished areas. Methods Ecol Evol 5:1061–1069

42. Magige FJ, Holmern T, Stokke S, Mlingwa C, Røskaft E (2009) Does illegal hunting affect density and behaviour of African grassland birds? A case study on ostrich (*Struthio camelus*). Biodivers Conserv 18:1361–1373. doi: 10.1007/s10531-008-9481-6

43. Marchand P, Garel M, Bourgoin G, Dubray D, Maillard D, Loison A (2014) Impacts of tourism and hunting on a large herbivore’s spatio-temporal behavior in and around a French protected area. Biol Conserv 177:1–11. doi: 10.1016/j.biocon.2014.05.022

44. Marsden SJ (1999) Estimation of parrot and hornbill densities using a point count distance sampling method. Ibis (Lond 1859) 141:327–390. doi: 10.1111/j.1474-919X.1999.tb04405.x

45. Maxwell SL, Fuller RA, Brooks TM, Watson JEM (2016) Biodiversity: The ravages of guns, nets and bulldozers. Nat 2016 5367615 536:143–145. doi: 10.1038/536143a

46. McGraw WS, Zuberbuhler K, Noe R (2007) Monkeys of the Taï Forest: An African primate community. Cambridge University Press, Cambridge

47. Miller DL, Burt ML, Rexstad EA, Thomas L (2013) Spatial models for distance sampling dataD: recent developments and future directions. Methods Ecol Evol 4:1001–1010. doi: 10.1111/2041-210X.12105

48. Miller DL, Rexstad E, Thomas L, Marshall L, Laake J (2016) Distance Sampling in R. bioRxiv 063891. doi: 10.1101/063891

49. Oates JF (2011) Primates of West Africa: A field guide and natural history. Conservation International, Bogotà

50. Ohashi H, Saito M, Horie R, et al. (2013) Differences in the activity pattern of the wild boar *Sus scrofa* related to human disturbance. Eur J Wildl Res 59:167–177. doi: 10.1007/s10344-012-0661-z

51. Palka DL, Hammond PS (2001) Accounting for responsive movement in line transect estimates of abundance. Can J Fish Aquat Sci 58:777–787

52. Papworth S, Milner-Gulland EJ, Slocombe K (2013) Hunted woolly monkeys (Lagothrix poeppigii) show threat-sensitive responses to human presence. PLoS One 8:e62000

53. Patten MA, Burger JC (2018) Reserves as double-edged sword: Avoidance behavior in an urban-adjacent wildland. Biol Conserv 218:233–239. doi: 10.1016/j.biocon.2017.12.033

54. Plumptre AJ, Sterling EJ, Buckland ST (2013) Primate census and survey techniques. Primate Ecol Conserv A Handb Tech 10–26

55. Railsback SF, Grimm V (2019) Agent-based and individual-based modeling: A practical introduction, second edition. Princeton University Press, Princeton

56. Römer H (2001) Ecological constraints for sound communication: from grasshoppers to elephants. In: Ecology of sensing. Springer, pp 59–77

57. Ruf T, Valencak T, Tataruch F, Arnold W (2006) Running speed in mammals increases with muscle n-6 polyunsaturated fatty acid content. PLoS One 1:e65

58. Sankey DWE, Storms RF, Musters RJ, Russell TW, Hemelrijk CK, Portugal SJ (2021) Absence of “selfish herd” dynamics in bird flocks under threat. Curr Biol 31, 3192–3198

59. Sheehan RL, Papworth S (2019) Human speech reduces pygmy marmoset (Cebuella pygmaea) feeding and resting at a Peruvian tourist site, with louder volumes decreasing visibility. Am J Primatol 81:e22967

60. Spaan D, Ramos-Fernández G, Schaffner CM, Pinacho-Guendulain B, Aureli F (2017) How survey design affects monkey counts: a case study on individually recognized spider monkeys (Ateles geoffroyi). Folia Primatol 88:409–420

61. Stokes EJ, Strindberg S, Bakabana PC, Elkan PW, Iyenguet FC, Madzoké B, Malanda GAF, Mowawa BS, Moukoumbou C, Ouakabadio FK, Rainey HJ (2010) Monitoring Great Ape and Elephant Abundance at Large Spatial Scales: Measuring Effectiveness of a Conservation Landscape. PLoS One 5:e10294. doi: 10.1371/JOURNAL.PONE.0010294

62. Suwanrat S, Ngoprasert D, Sutherland C, Suwanwaree P, Savini T (2015) Estimating density of secretive terrestrial birds (Siamese Fireback) in pristine and degraded forest using camera traps and distance sampling. Glob Ecol Conserv 3:596–606. doi: 10.1016/j.gecco.2015.01.010

63. Tarakini T, Crosmary WG, Fritz H, Mundy P (2014) Flight behavioural responses to sport hunting by two African herbivores. South African J Wildl Res 44:76–83. doi: 10.3957/056.044.0110

64. Thomas L, Buckland ST, Rexstad EA, Laake JL, Strindberg S, Hedley SL, Bishop JRB, Marques TA, Burnham KP (2010) Distance software: Design and analysis of distance sampling surveys for estimating population size. J Appl Ecol 47:5–14. doi: 10.1111/j.1365-2664.2009.01737.x

65. Tuomainen U, Candolin U (2011) Behavioural responses to human-induced environmental change. Biol Rev 86:640–657. doi: 10.1111/j.1469-185X.2010.00164.x

66. Vabø R, Nøttestad L (1997) An individual based model of fish school reactions: Predicting antipredator behaviour as observed in nature. Fish Oceanogr 6:155–171. doi: 10.1046/j.1365-2419.1997.00037.x

67. Watanabe K (1981) Variations in group composition and population density of the two sympatric Mentawaian leaf-monkeys. Primates 22:145–160. doi: 10.1007/BF02382606

68. White JW, Rassweiler A, Samhouri JF, Stier AC, White C (2014) Ecologists should not use statistical significance tests to interpret simulation model results. Oikos 123:385–388

69. Whittaker D, Knight RL (1998) Understanding wildlife responses to humans. Wildl Soc Bull 26:312–317

